# Sparse recurrent excitatory connectivity in the microcircuit of the adult mouse and human cortex

**DOI:** 10.1101/292706

**Authors:** Stephanie C. Seeman, Luke Campagnola, Pasha A. Davoudian, Alex Hoggarth, Travis A. Hage, Alice Bosma-Moody, Christopher A. Baker, Jung Hoon Lee, Stefan Mihalas, Corinne Teeter, Andrew L. Ko, Jeffrey G. Ojemann, Ryder P. Gwinn, Daniel L. Silbergeld, Charles Cobbs, John Phillips, Ed Lein, Gabe J. Murphy, Christof Koch, Hongkui Zeng, Tim Jarsky

**Affiliations:** Allen Institute for Brain Science, Seattle, WA; Regional Epilepsy Center at Harborview Medical Center, Seattle, WA; Department of Neurological Surgery, University of Washington School of Medicine, Seattle, WA; Epilepsy Surgery and Functional Neurosurgery, Swedish Neuroscience Institute, Seattle, WA; The Ben and Catherine Ivy Center for Advanced Brain Tumor Treatment, Swedish Neuroscience Institute, Seattle, WA

## Abstract

Generating a comprehensive description of cortical networks requires a large-scale, systematic approach. To that end, the Allen Institute is engaged in a pipeline project using multipatch electrophysiology, supplemented with 2-photon optogenetics, to characterize connectivity and synaptic signaling between classes of neurons in adult mouse and human cortex. We focus on producing results detailed enough for the generation of computational models and enabling comparison with future studies. Here we report our examination of intralaminar connectivity within each of several classes of excitatory neurons. We find that connections are sparse but present among all excitatory cell types and layers we sampled, with the most sparse connections in layers 5 and 6. Almost all mouse synapses exhibited short-term depression with similar dynamics. Synaptic signaling between a subset of layer 2/3 neurons; however, exhibited facilitation. These results contribute to a body of evidence describing recurrent excitatory connectivity as a conserved feature of cortical microcircuits.

## Introduction

Generating well-informed, concrete, testable hypotheses about how the cortex represents and processes information requires experimental efforts to characterize the connectivity and dynamics of cortical circuit elements as well as efforts to build models that integrate results across studies (Sejnowski et al. 1988). Estimates of connectivity and synaptic properties vary widely between experiments due to differences in model organisms, experimental parameters, and analytic methods. This variability limits our ability to generate accurate, integrative computational models.

Addressing this problem requires standardized experimental methods and large-scale data collection in order to characterize synaptic connections between the large number of potential cell types (Tasic et al. 2016). Although it may be possible to infer part of these results based solely on anatomical constraints (Markram et al. 2015), evidence has shown that the rate of connectivity and properties of synaptic signals can depend on the identity of the pre-and postsynaptic neuron (Reyes et al. 1998, Galarreta and Hestrin, 1998). To collect standardized data at scale, the Allen Institute for Brain Science has built a pipeline to characterize local, functional connectivity in the adult mouse and human cortex. Here we report on the characteristics of recurrent, intralayer connectivity among pyramidal neurons generated during the pipeline’s system integration test—an end-to-end test of the pipeline’s hardware, software, and workflow for a subset of all synaptic connections.

Recurrent excitatory connectivity is a common feature in computational models of cortical working memory, receptive field shaping, attractor dynamics, and sequence storage (Camperi & Wang 1998, Olshausen & Field 1996, Brunel et al. 2016, Pernice et al. 2018). Empirical measurements of recurrent connectivity and synaptic properties are needed in order to constrain and validate these models. However, characterizing recurrent connectivity in a standardized, high throughput manner is challenging because the synaptic connections can be sparse and weak (Braitenberg and Schüz 1998, Lefort 2009). Furthermore, most measurements of recurrent connectivity have been performed in juvenile rodents, leading to a recent debate over the rate of connectivity in the adult cortex (Biane et al. 2015, Barth et al. 2016, Jiang et al. 2016).

The data reported here demonstrate that sparse recurrent connectivity is present among excitatory neurons in all layers of adult mouse and human cortex. Using a novel automated method for systematically estimating connectivity across experiments we further demonstrate that different populations of adult mouse pyramidal neurons exhibit characteristic distance-dependent connectivity profiles and short-term dynamics. We quantify and compare differences in short-term dynamics with a mechanistic computational model.

## Results

Here, we report on data collected during the system integration test of a new Allen Institute for Brain Science pipeline for the systematic characterization of local connectivity in the adult cortex. We performed *in vitro* whole-cell recordings from up to eight excitatory neurons simultaneously. To assess connectivity, trains of action potentials were evoked in each cell, one at a time, while recording current clamp synaptic responses in all other cells. We probed 2324 putative connections (200 minimum per layer) in mouse V1 and 332 putative connections (35 minimum per layer) in human frontal and temporal cortex (Table 1). Connections were identified by the presence of excitatory PSPs (EPSPs) evoked with a short latency and low jitter, following the presynaptic spike, consistent with a monosynaptic connection (Figure S1). Recurrent connectivity was observed in layer 2/3 through layer 6 of mouse primary visual cortex and layer 2 through layer 5 of the human cortex.

**Table 1:**
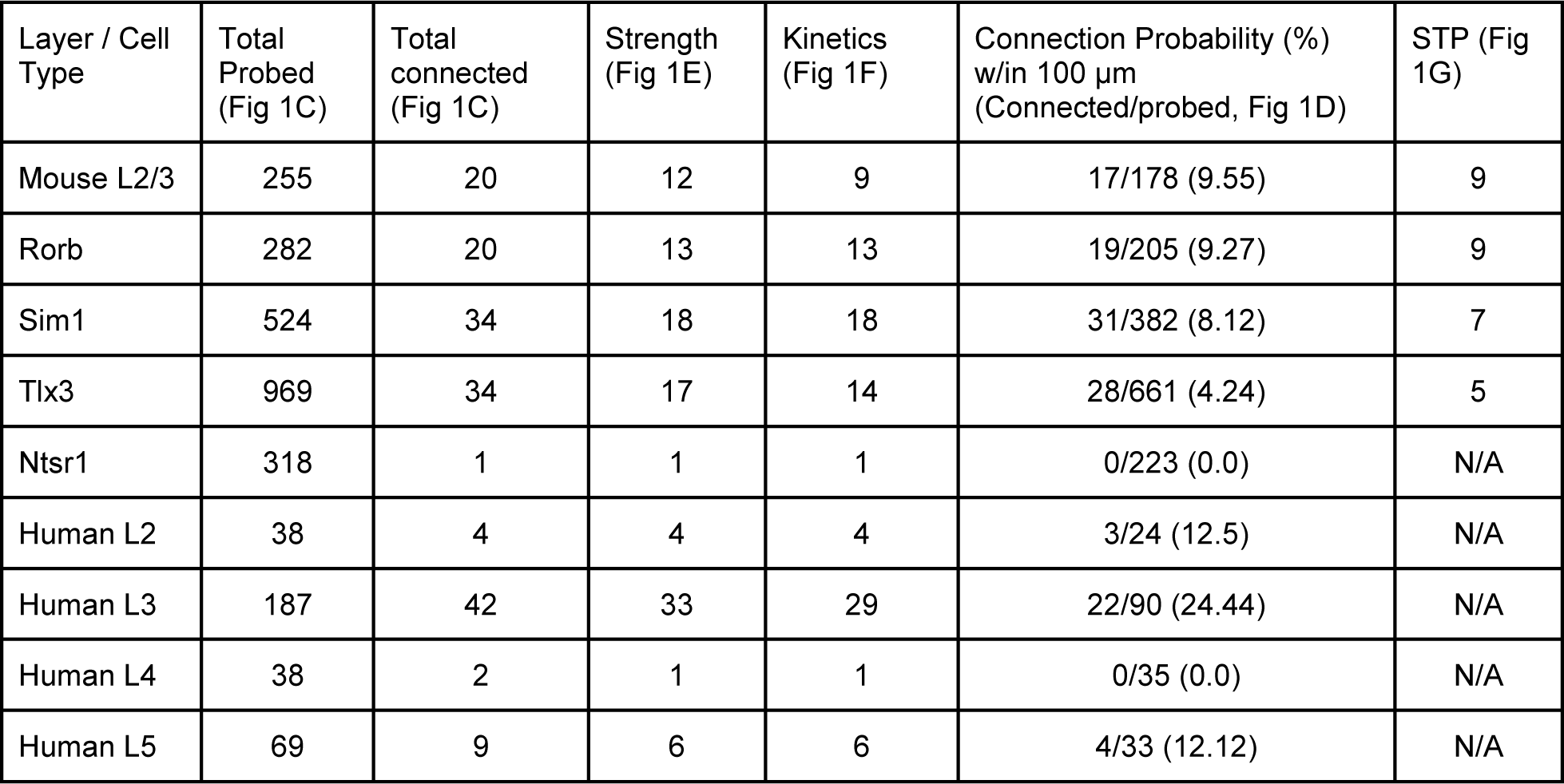
Number of potential connections used in each connection analysis.

| Layer / Cell Type | Total Probed (Fig 1C) | Total connected (Fig 1C) | Strength (Fig 1E) | Kinetics (Fig 1F) | Connection Probability (%) w/in 100 μm (Connected/probed, Fig 1D) | STP (Fig 1G) |
| --- | --- | --- | --- | --- | --- | --- |
| Mouse L2/3 | 255 | 20 | 12 | 9 | 17/178 (9.55) | 9 |
| Rorb | 282 | 20 | 13 | 13 | 19/205 (9.27) | 9 |
| Sim1 | 524 | 34 | 18 | 18 | 31/382 (8.12) | 7 |
| Tlx3 | 969 | 34 | 17 | 14 | 28/661 (4.24) | 5 |
| Ntsr1 | 318 | 1 | 1 | 1 | 0/223 (0.0) | N/A |
| Human L2 | 38 | 4 | 4 | 4 | 3/24 (12.5) | N/A |
| Human L3 | 187 | 42 | 33 | 29 | 22/90 (24.44) | N/A |
| Human L4 | 38 | 2 | 1 | 1 | 0/35 (0.0) | N/A |
| Human L5 | 69 | 9 | 6 | 6 | 4/33 (12.12) | N/A |

### Properties of intralaminar excitatory synaptic signaling in mouse cortex

Layer and projection-specific subclasses of excitatory neuron populations were identified either by *post-hoc* morphologic evaluation in layer 2/3 or transgenic labelling to target layers 4-6 (Rorb, Tlx3, Sim1, and Ntsr1, respectively; Figure 1A). Layer 5 recordings were subdivided into subcortical projecting cells (Sim1) or corticocortical projecting cells (Tlx3). We probed 2384 potential connections (layer 2/3: 255, Rorb: 282, Sim1: 524, Tlx3: 969, Ntsr1: 318) across these excitatory populations in mouse cortex (Table 1). Connections were detected between 107 putative pre-and post-synaptic partners (layer 2/3: 20, Rorb: 20, Sim1: 34, Tlx3: 34, Ntsr1: 1; Table 1). For >75% of the recorded cells, we recovered a biocytin fill (Fig 1A) and for all cells we obtained an epifluorescent image stack (Fig 1B).

**Figure 1.**
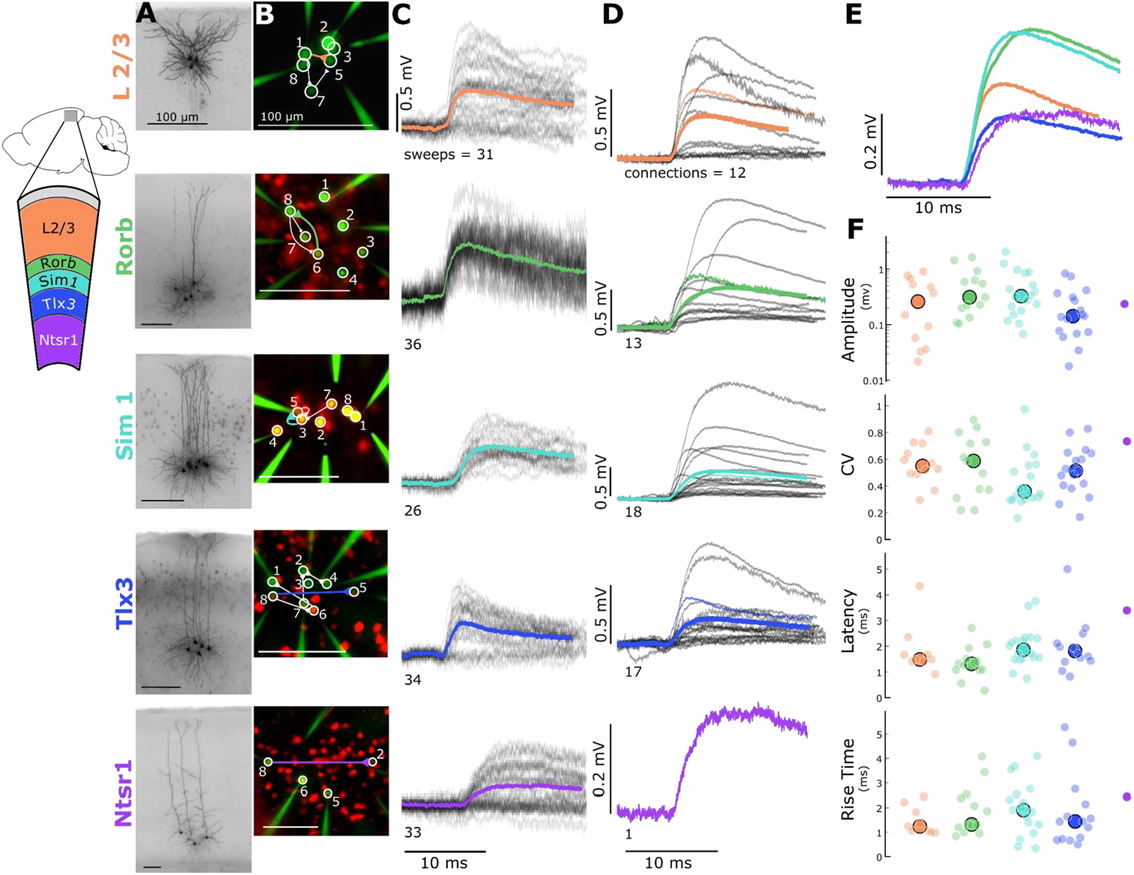
Electrophysiological recordings of evoked excitatory synaptic responses between individual cortical pyramidal neurons in mouse primary visual cortex. **A**. Cartoon illustrating color, Cre-type, and cortical layer mapping in slice recording region (V1). Example maximum intensity projection images of biocytin-filled pyramidal neurons for L2/3 and each Cre-type. **B**. Example epifluorescent images of neurons showing Cre-dependent reporter expression and/or dye-filled recording pipettes. Connection map is overlaid on the epifluorescent image (colored: example connection shown in C). **C**. Spike time aligned EPSPs induced by the first AP of all ≤ 50 Hz stimulus trains for a single example connection (individual pulse-response trials: grey; average: colored). **D**. First pulse average, like in C., for all connections within the synaptic type; grey: individual connections; thin-colored: connection highlighted in C; thick-colored: grand average of all connections. **E**. Overlay of grand average for each connection type. F. EPSP amplitude (in log units), CV of amplitude, latency, and rise time of first pulse response for each Cre type (small circles) with the grand median (large).

In layers 2/3 through 5 we were able to sample sufficiently to characterize the strength and kinetics in recurrent connections. To measure these features with minimal influence of STP, only the first response on each sweep (ITI = 15s) was included for this analysis. A subset of connections from each group were chosen based on quality control metrics and inclusion criteria (see methods; Fig S1). Figure 1C shows EPSPs recorded from one example connection found in each of the five chosen excitatory cell groups. For the large majority of connections, it was not possible to unequivocally distinguish synaptic failures from detection failures, thus we used the mean response from all sweeps (Fig 1C) to evaluate the EPSP features.

Consistent with previous reports that recurrent connectivity is weak (Lefort 2009), we found that a majority of the connections had amplitudes less than 0.5 mV. In this small sample we did not observe statistical difference in the EPSP amplitudes (Fig 1E,F) between groups (KW p = 0.07) although the median Rorb EPSP (0.312 ± 0.485 mV) was more than 2 times larger than the median Tlx3 EPSP (0.142 ± 0.231 mV). This was likely driven by the heterogeneity in single pulse responses within groups; the range of amplitudes for layer 2/3 (0.032 - 0.902 mV), Rorb (0.105 – 1.626 mV), Sim1 (0.068 - 1.254 mV), and Tlx3 (0.02 - 0.833 mV) spanned an order of magnitude (we could not assess the range of recurrent Rorb or Ntsr1 connections due to the low number of connections measured). The mean EPSP amplitude was consistently larger than the median (Table 2), likely due to a skewed (long-tailed) distribution of response amplitudes. A similar observation in the rat visual cortex, and mouse somatosensory cortex, has led to the suggestion that rare, large-amplitude connections are important for reliable information processing (Song et al 2005, Lefort 2009, Cossell et al. 2015). The majority of EPSP latencies were less than 2.5 ms, and similar across populations (KW p= 0.17), consistent with a direct, monosynaptic connection between recorded neurons.

**Table 2:** Mouse single-pulse response properties (median, except where indicated)

| Connection Type/Feature | L2/3 | Rorb | Sim1 | Tlx3 |
| --- | --- | --- | --- | --- |
| Amplitude (mV ±SD) | $0.260 \pm 0.306$ | $0.312 \pm 0.485$ | $0.326 \pm 0.499$ | $0.142 \pm 0.231$ |
| Amplitude mean (mV ±SD) | $0.342 \pm 0.092$ | $0.544 \pm 0.140$ | $0.520 \pm 0.121$ | $0.236 \pm 0.058$ |
| Latency (ms ±SD) | $1.484 \pm 0.945$ | $1.313 \pm 0.604$ | $1.861 \pm 0.745$ | $1.812 \pm 0.971$ |
| Rise Time (ms ±SD) | $1.240 \pm 0.528$ | $1.320 \pm 0.967$ | $1.905 \pm 1.046$ | $1.440 \pm 1.425$ |
| CV (±SD) | $0.549 \pm 0.142$ | $0.587 \pm 0.228$ | $0.358 \pm 0.189$ | $0.513 \pm 0.174$ |

We could not directly quantify synaptic failures and thus calculated the coefficient of variation of synaptic amplitudes (CV; Fig 1F) to assess release probability. The CV of each connection describes the variability in a particular response in relation to the mean (ratio of standard deviation to mean) and is negatively correlated with release probability (Markram et al, 1997). The range of coefficient of variation in our data suggests differences in release probability across cells as well as between cell types (KW p=0.02). This is consistent with STP modeling results (see Fig 6).

**Figure 6.**
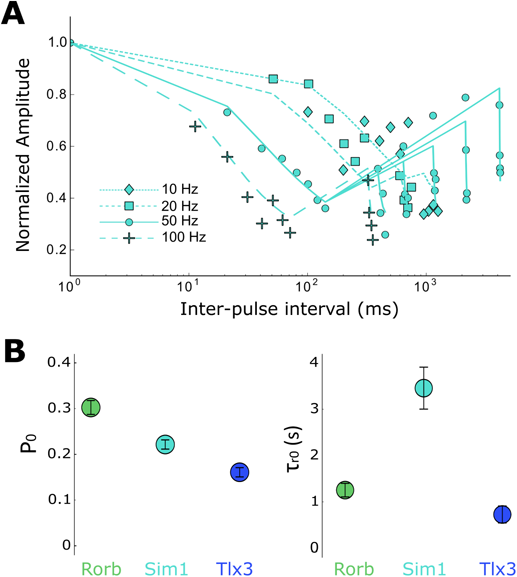
Modeling of short-term depression in recurrent Rorb, Sim1, and Tlx3 connections. **A**. Sim1 Cre average dynamic response; Same data as in Fig 5C, top plotted on a log-X time scale with modeling fits overlaid. **B**. Results of model for parameters P0 and *τ*_r0_. Values are means with standard error of the covariance matrix. Paired Z-scores (Eq. 6) in Table 4.

### Properties of intralaminar excitatory synaptic signaling in human cortex

To what extent is recurrent connectivity in mouse V1 representative of connectivity in other regions and species? To make this comparison, we performed multipatch recordings in human surgical specimens from temporal and frontal cortex. We sampled recurrent intralayer connectivity in all layers containing pyramidal cells. Excitatory cells were identified either by the polarity of the synaptic currents evoked by stimulating the cell or by their morphology visualized via biocytin (Fig 2A) or fluorescent dye (Fig 2B). We found 4 connections between layer 2 pyramidal cells (38 probed), 42 connections between layer 3 pyramidal cells (187 probed), 2 connections between layer 4 pyramidal cells (38 probed), and 9 connections between layer 5 pyramidal cells (69 probed). We did find one connection in layer 6 but have not probed this layer sufficiently to say more (2 probed connections). Human cortex had a higher connectivity rate and mean amplitude (Fig 2C, D) compared to mouse cortex (despite a higher [Ca]_e_ in mouse), consistent with previous reports (Molnar et. al 2008). Layers 2, 3, and 5 had a sufficient number of connections to characterize strength and kinetics however, we found no significant differences in response properties between layers (amplitude p=0.51, latency p=0.24, rise time =0.13, Table 3). We did observe differences in CV between layers 2, 3, and 5 (p=0.03, Table 3) suggesting layer-specific differences in release probability of recurrent connections, similar to findings in mouse V1.

**Table 3:** Human single-pulse response properties (median, except where indicated)

| Connection Type/Feature | L2 | L3 | L5 |
| --- | --- | --- | --- |
| Amplitude (mV ±SD) | 0.280 ± 0.092 | 0.342 ± 0.660 | 0.615 ± 0.632 |
| Amplitude mean (mV ±SD) | 0.301 ± 0.092 | 0.549 ± 0.660 | 0.835 ± 0.632 |
| Latency (ms ±SD) | 0.871 ± 0.764 | 1.555 ± 0.989 | 1.639 ± 0.542 |
| Rise Time (ms ±SD) | 1.765 ± 0.552 | 1.600 ± 1.180 | 1.155 ± 0.419 |
| CV (±SD) | 0.740 ± 0.267 | 0.393 ± 0.228 | 0.290 ± 0.092 |

**Table 4:**
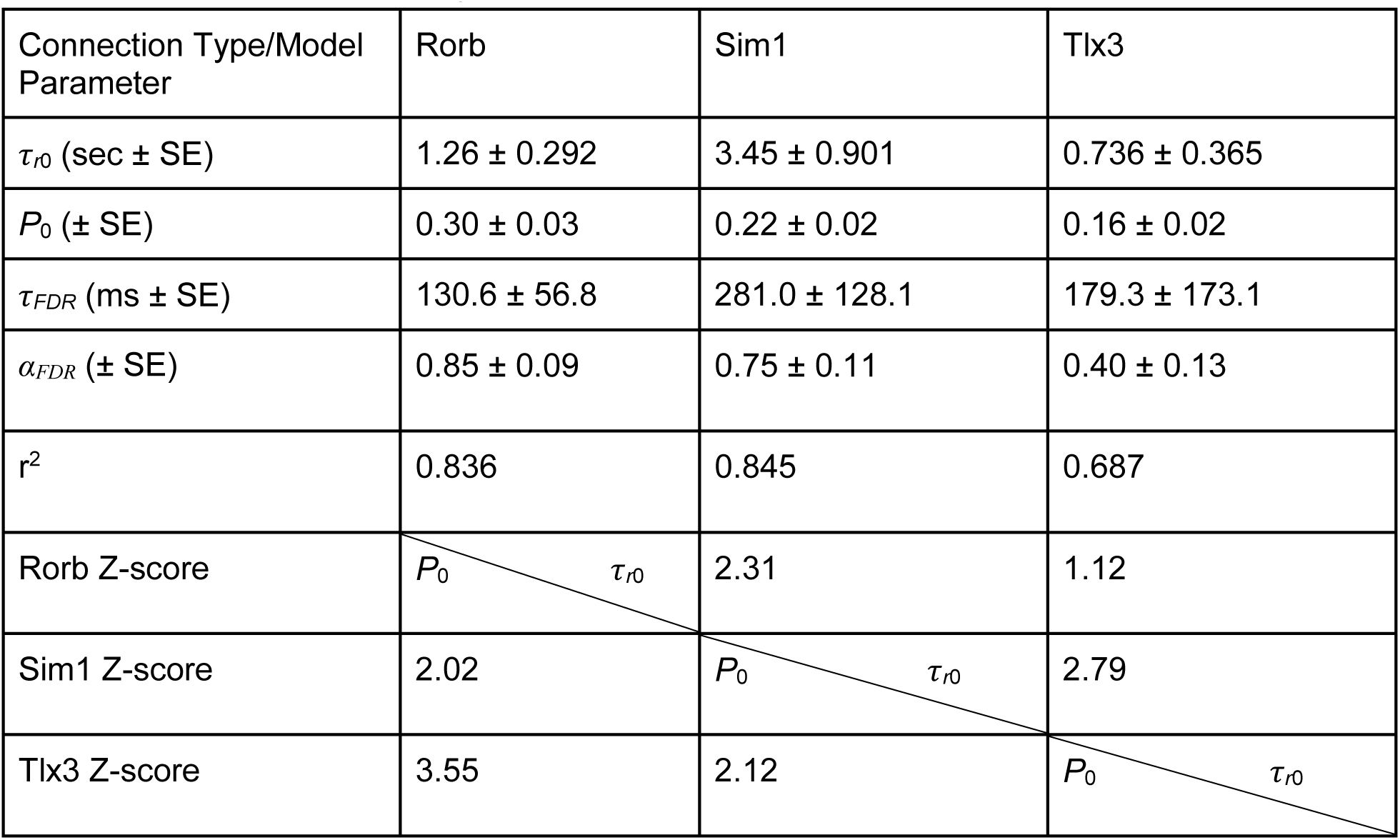
Model of short-term dynamics.

| Connection Type/Model Parameter | Rorb | Sim1 | Tlx3 |
| --- | --- | --- | --- |
| $\tau_{r0}$ (sec ± SE) | $1.26 \pm 0.292$ | $3.45 \pm 0.901$ | $0.736 \pm 0.365$ |
| $P_0$ (± SE) | $0.30 \pm 0.03$ | $0.22 \pm 0.02$ | $0.16 \pm 0.02$ |
| $\tau_{FDR}$ (ms ± SE) | $130.6 \pm 56.8$ | $281.0 \pm 128.1$ | $179.3 \pm 173.1$ |
| $\alpha_{FDR}$ (± SE) | $0.85 \pm 0.09$ | $0.75 \pm 0.11$ | $0.40 \pm 0.13$ |
| $r^2$ | 0.836 | 0.845 | 0.687 |
| Rorb Z-score | $P_0$ $\tau_{r0}$ | 2.31 | 1.12 |
| Sim1 Z-score | 2.02 | $P_0$ $\tau_{r0}$ | 2.79 |
| Tlx3 Z-score | 3.55 | 2.12 | $P_0$ $\tau_{r0}$ |

**Figure 2.**
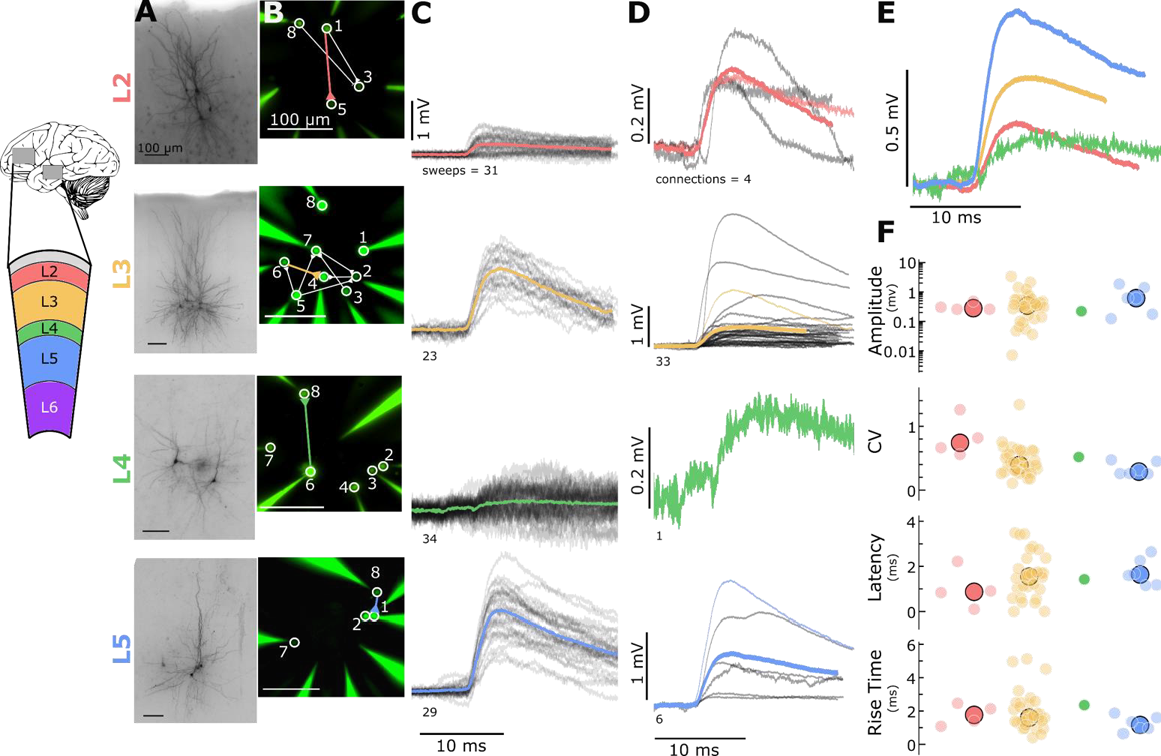
Electrophysiological recordings of evoked excitatory synaptic responses between individual human cortical pyramidal neurons. **A**. Cartoon illustrating color and cortical layer mapping in slice recording region (temporal or frontal cortex). Example maximum intensity projection images of biocytin-filled pyramidal neurons for layers 2-5. **B**. Example epifluorescent images of neurons showing dye-filled neurons and recording pipettes. Connection map is overlaid on the epifluorescent image (colored: example connection shown in C). **C**. Spike time aligned EPSPs induced by the first AP of all ≤ 50 Hz stimulus trains for a single example connection (individual pulse-response trials: grey; average: colored). **D**. First pulse average, like in C., for all connections within the synaptic type; grey: individual connections; thin-colored: connection highlighted in C; thick-colored: grand average of all connections. **E**. Overlay of grand average for each connection type. **F**. EPSP amplitude, CV of amplitude, latency, and rise time of first pulse response for each layer (small circles) with the grand mean (large circles).

It is reasonable to question if the recurrent connections we see in the human are related to the disease etiology. Although we cannot rule this out, we saw no significant differences in overall connectivity between tumor and epilepsy derived specimens (p = 0.833, Fisher’s Exact Test). We also found recurrent connections across multiple cortical regions and disease states in the human. Taken together, this may indicate that our results capture a common architecture of the mouse and human microcircuit.

#### Detection limit of synaptic responses

When using whole cell recordings to characterize synaptic connectivity, a major limitation is that some synaptic currents may be too weak to be detected at the postsynaptic soma. Detection limits are influenced by the kinetics of PSPs, the duration of the experiment (number of averaged pulse-responses), and the amplitude and kinetics of background noise, including the properties of spontaneous PSPs. Worse, we lack even a basic estimate of the fraction of synapses that could be missed. For this reason, measurements of connectivity from multipatch experiments should be considered lower-bound estimates. Additionally, apparent connectivity may be systematically biased by cell type differences in synaptic strength, making it more difficult to interpret any measured differences in connectivity.

To address these issues, we simulated EPSPs of varying, known strength, and used a machine classifier to measure the probability that synapses could escape detection (see Methods). For each putative connection probed, we estimated the minimum PSP amplitude that could be detected. When compared to the measured amplitudes of detected PSPs, it is apparent that our data set does contain false negatives (Fig 3A, area under red dashed line). The classifier was trained to detect connections based on features previously extracted from the averaged pulse response (Fig 3B) and from the distributions of features measured on individual pulse responses (Fig 3C). Using background recording data from the postsynaptic cell, we then generated several sets of artificial pulse responses and measured the probability that the classifier would detect these, while varying the rise time and mean amplitude of the PSPs (Fig 3D).

**Figure 3.**
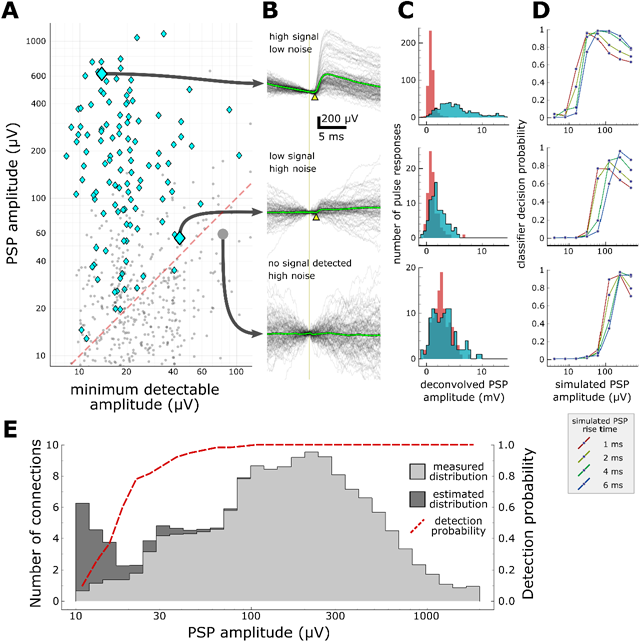
Characterization of synapse detection limits. **A**. Scatter plot showing measured PSP amplitude versus minimum detectable amplitude for each tested pair. Detected synapses (manually annotated) are shown as blue diamonds; pairs with no detected synaptic currents are grey dots. The region under the red dashed line denotes the region in which synaptic connections are likely to be misclassified as unconnected. Three example putative connections are highlighted in A and described further in panels B-D. One connection (top row) was selected for its large amplitude PSP and low background noise. Another connection (middle row) is harder to detect (PSP onset marked by yellow arrowhead) due to low amplitude and high background noise. The bottom row shows a recorded pair that was classified as unconnected. **B**. A selection of postsynaptic current clamp recordings in response to presynaptic spikes. Each row contains recordings from a single tested pair. The vertical line indicates the time of presynaptic spikes, measured as the point of maximum dV/dt in the presynaptic recording. Yellow triangles indicate the onset of the PSP. **C**. Histograms showing distributions of peak response values measured from deconvolved traces (see Methods). Red area indicates measurements made on background noise; blue area indicates measurements made immediately following a presynaptic spike. **D**. Characterization of detection limits for each example. Plots show the probability that simulated PSPs would be detected by a classifier, as a function of the rise time and mean amplitude of the PSPs. Each example has a different characteristic detection limit that depends on the recording background noise and the length of the experiment. **E**. An estimate of the total number of false negatives across the entire dataset. The measured distribution of PSP amplitudes is shown in light grey (smoothed with a gaussian filter with σ=1 bin). The estimated correction show in dark grey is derived by dividing the measured distribution by the overall probability of detecting a synapse (red dashed line) at each amplitude.

With this approach, we can place constraints for any given experiment on the properties of putative missed synaptic connections. Recordings with low background noise and adequate averaging will generally allow the detection of very small synaptic currents (Fig 3D, top panel has a detection limit 10-20 µV), whereas lower quality recordings will have higher detection thresholds and will report lower connectivity rates (Fig 3D, bottom panel has a poor detection limit near 100 µV). Likewise, PSPs with shorter rise time (or other properties that distinguish the PSP from background) are more likely to be detected (Fig 3D). These results suggest that the differences in experimental protocol between studies (for example, the amount of averaging done for each connection) can have a substantial impact on the apparent connectivity reported, but also that future studies could reconcile these differences by carefully characterizing their detection limits.

The results in Figure 3A suggest another tantalizing opportunity: if we know the area that contains false negatives (under the red dashed line) and the density of true positives above the line, then we can make a first-order estimate of the number of connections that were missed across a series of experiments. Figure 3E shows the distribution of PSP amplitudes across all detected synapses (light grey area) as well as the curve representing the probability that synapses would be detected at any amplitude. Dividing the measured distribution by the probability of detection yields a corrected distribution (dark grey) with an overall 10% increase in connectivity. Although it is clear that this estimate becomes unstable as the detection probability approaches zero, a surprising result of this analysis is that, for moderate amplitudes of 20-100 µV, we expect to see very few false negatives. Depending on the expected prevalence of weaker connections < 20 µV, this hints that our current sampling methods may be adequate to detect the vast majority of synaptic connections. We should be cautious, however, in our interpretation of this result--the analysis relies on several assumptions about the behavior of the classifier and the realism of the simulated PSPs. Ultimately, the approach must be validated against a larger dataset.

### Connection probability of excitatory synapses

Estimates of connectivity vary widely across studies, in part due to methodological differences. In addition to the effects of detection sensitivity described above, estimated connection probability is affected by the intersomatic distances over which connections are sampled. This spatial distribution of connections may also offer insight into the organization of functional microcircuits. In mouse, connectivity in layer 2/3, Rorb, and Sim1 neurons within 100 µm was 201 similar (∼10%; Fig 4, left; connected/probed, L2/3: 17/178, Rorb: 19/205, Sim1: 49/507). However, within this range, Tlx3 connectivity was markedly lower (∼4%; Tlx3: 32/818,). Ntsr1 connectivity was significantly more sparse as only one connection was detected (out of 312 probed) at a distance of 163 µm (p<0.05 relative to all other groups). Most connectivity versus distance profiles (Fig 4B) showed a progressive reduction in the connection probability with increasing distance.

**Figure 4.**
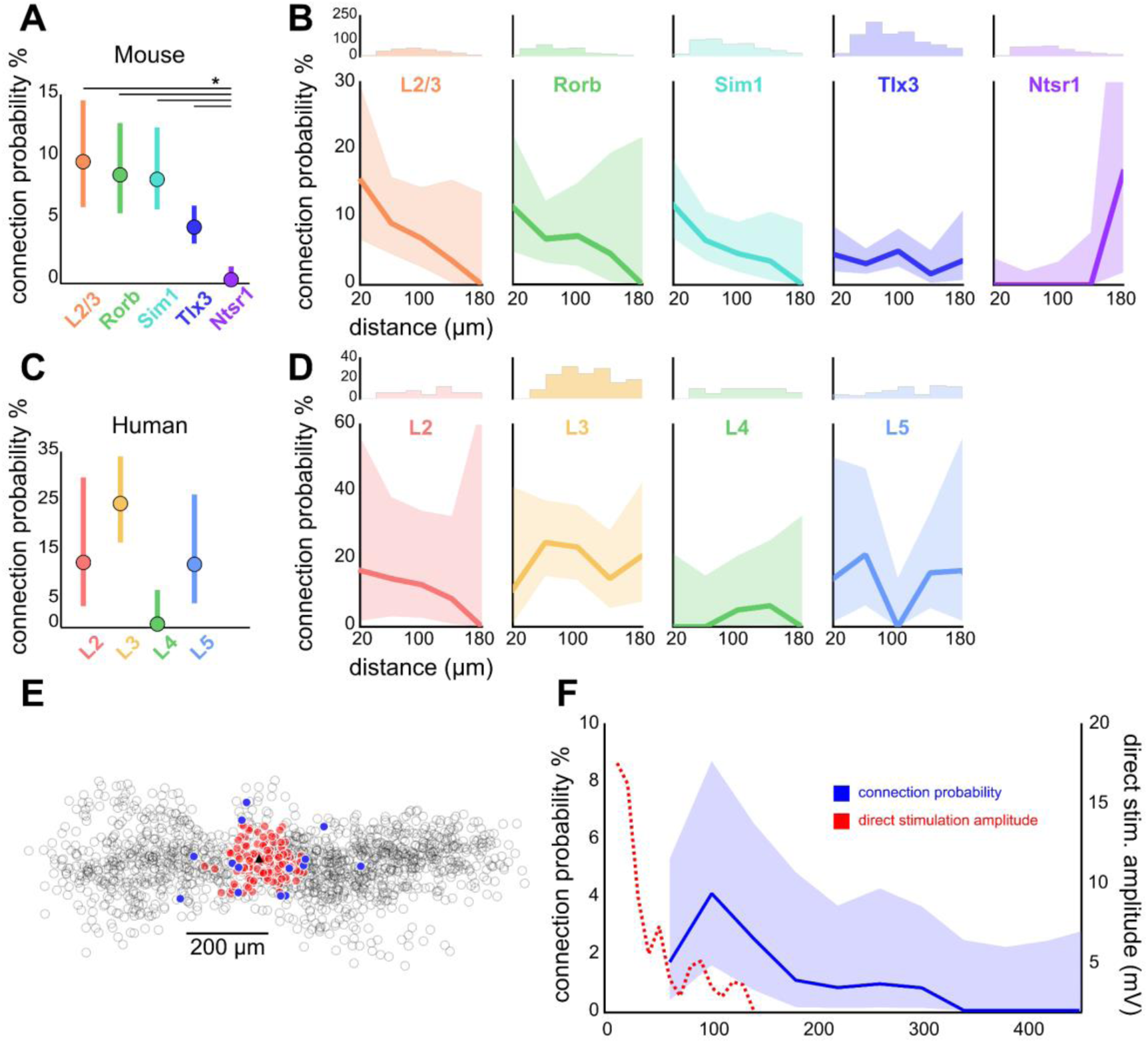
Distance dependent connectivity profiles of mouse and human E-E connections. **A**. Recurrent connection probability and distribution of connections for mouse Cre lines and layer 2/3. Mean connection probability (filled circles) and 95% confidence intervals (bars) for connections probed within 100 µm. Asterisk (*) indicates p-value <0.05 (Bonferroni corrected). **B**. Connection probability over distance for mouse Cre lines and layer 2/3. *Top*: Histogram of putative connections probed. *Bottom:* Mean connection probability (thick line) with 95% confidence intervals (shading). **C**. Like-to-like connection probability and distribution of connections between human pyramidal neurons. Mean connection probability (filled circles) and 95% confidence intervals (bars) for connections probed within 100 µm. **D**. Connection probability over distance for human pyramidal neurons, formatted as in panel B. **E**. Tlx3-Tlx3 connection probability measured by two-photon mapping. X-Y distance distribution of connections probed onto a postsynaptic cell (black triangle), detected presynaptic neurons (filled circles), no connection detected (empty circles), and direct event artifact due to undesired activation of opsin in the dendritic arbor of the recorded cell (red circles). **F**. Connection probability and stimulation artifact over distance measured by two-photon mapping. Mean connection probability vs. distance (blue line; starting at 50 µm) with 95% confidence intervals (shading) and direct event artifact amplitude vs. distance (dotted red line) for Tlx3-Tlx3 connections probed with two-photon stimulation.

Utilizing a multipatch technique limits our ability to probe connectivity at high density and far distances. Two-photon optogenetic stimulation, which allows for focal stimulation of many (mean=57 cells, range=8-117) presynaptic cells in a single experiment, and critically, enables the intersomatic distance between those cells to be greater than is generally feasible with multipatch experiments, was used to overcome these limitations. ReaChR expressing Tlx3-Cre neurons in layer 5 were photo-stimulated while one or two putative postsynaptic cells were monitored in whole-cell current clamp configuration (Fig 4E). With this technique, using 9 mice, we found a similar connection probability over the distance range of multipatch experiments (4/167, 2.40%) and reduced connectivity at extended distances up to 785 µm (Fig 4E; 10/1290, 0.78%) with the furthest connection found at 300 µm.

Overall, connectivity in human cortex was significantly higher than that in mouse (human 19%, mouse 6.8%, Fisher’s p<0.001). In human cortex, layer 2 and layer 5 connectivity showed similar connection probability (∼12%; layer 2: 3/24; layer 5: 4/33; Fig 4C), while layer 3 recurrent connectivity was approximately double (24%, 22/90). More data is needed to accurately resolve the distance dependence of recurrent connectivity in the human (Fig 4D).

### Short-term plasticity of excitatory synapses

For a subset of synaptic connections in mouse cortex (Fig S1G), we characterized the short-term synaptic dynamics. We probed short-term dynamics with stimulus trains consisting of eight pulses to induce STP, followed by a delay and four more pulses to measure recovery (Fig 5A, left). The eight initial pulses allowed responses to reach a steady state, from which we could characterize the extent of depression (or facilitation) at frequencies from 10 - 100 Hz. The 50 Hz stimulation protocol had additional recovery intervals ranging from 250 - 4000 ms (Fig 5A, right). Figure 5B shows average synaptic responses to a 50 Hz stimulus with 8 initial pulses followed by 4 pulses at a 250 ms delay from individual Sim1-Sim1 connections shown in grey, and the grand average overlaid (cyan). We used exponential deconvolution (Fig 5B, middle; Eq. 2) to estimate the amplitudes of individual PSPs in the absence of temporal summation (arising from the relatively long cell membrane time constant).

**Figure 5.**
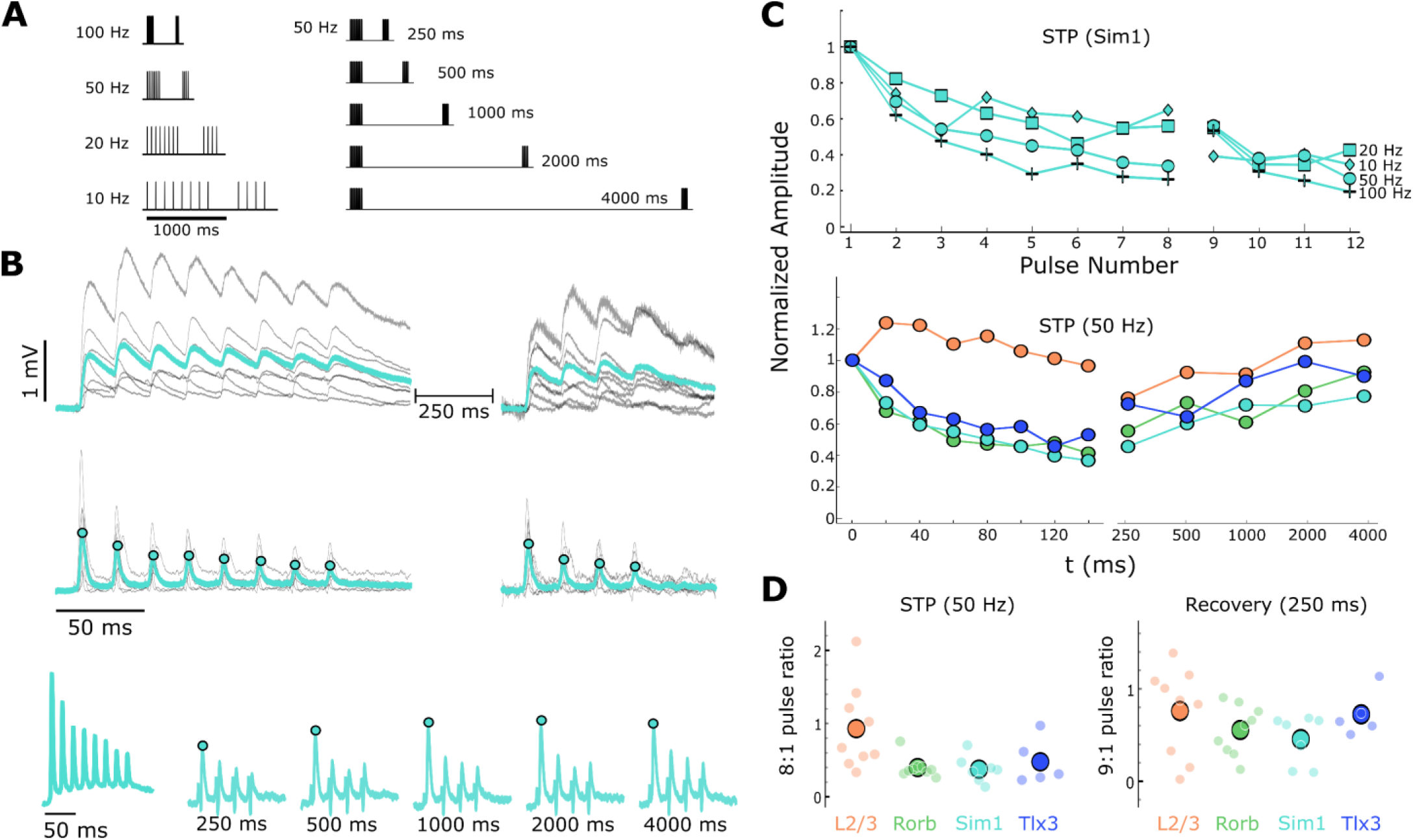
Short-term dynamics of mouse recurrent connections by Cre-type and layer. **A**. Schematic of STP and STP recovery stimuli. **B**. Sim1-Cre EPSPs in response to a 50 Hz stimulus train (top; 8 induction pulses and 4 recovery pulses delayed 250 ms; individual connection: gray traces; blue: Sim1-Cre average EPSP at 50Hz). Exponential deconvolution followed by lowpass filter of EPSPs above (middle, filled circles: pulse amplitudes in C.). Exponential deconvolution of 50 Hz stimulus with all 5 recovery time points in A (bottom, filled circles: pulse amplitudes in C.). **C**. The mean normalized amplitude of deconvolved response versus pulse number at multiple stimulation frequencies for Sim1-Cre (top). Normalized amplitude of the deconvolved response at 50 Hz with first recovery pulse at each interval for each Cre-type and L2/3 connections (bottom). **D**. The depth of depression during 50Hz induction (left) as measured by the amplitude ratio of the 8th to 1st pulse for each Cre-type and layer (small circles) and grand mean (large circles). Amount of recovery at 250 ms latency (right) for each Cre-type and layer (small circles) and grand mean (large circles).

We measured the peak amplitude of the deconvolved response for every pulse (cyan dots) and normalized to the first pulse in the train in order to characterize short-term dynamics across four frequencies. Fig 5C (top) highlights frequency dependent depression in recurrent Sim1 connections. We similarly measured recovery from short-term effects at various time delays (Fig 5B, bottom, cyan dots) for layer 2/3, Rorb, Sim1, and Tlx3 connections (Fig 5C, bottom). A simple analysis of the magnitude of, and recovery from, short-term plasticity is shown in Fig 5D. The amplitude ratio of the last pre-recovery interval pulse (8) to the first shows that layer 2/3 exhibits less depression than Rorb, Sim1, and Tlx3 (Fig 5D, left). All types showed a similar time-course of recovery as measured by the ratio of the first recovery pulse (9) to the first induction pulse (Fig 5D, right). Recovery also tended to be rapid in the first 250 ms followed by a slower, incomplete recovery measured at 4 seconds (Fig 5C, middle).

To better capture the dynamic processes contributing to short-term plasticity and how they may differ among connection types we turned to a model of short-term dynamics which has been well described (Hennig, 2013; Mongillo et al., 2008; Richardson et al., 2005). Rorb, Sim1, and Tlx3 synapses were modelled with depression (Eq. 4, methods) and use-dependent replenishment (Eq. 5, methods). This model performed well (Fig 6A, Sim1 connections r^2^ = 0.845, Table 4) in capturing depression during the 8 initial pulses at various frequencies (Fig 6A) as well as modeling recovery at various delays (Fig 6A, circles). From this model we can extract free parameters such as P_0_ which describes release probability and *τ*_r0_which describes the time course of recovery from depression. Rorb connections had the largest release probability (P_0_ = 0.30, Fig 6B, left) which is consistent with faster entry into depression (Fig 5C, bottom pulse 2). Conversely, Tlx3 connections had a lower initial release probability (P0 = 0.16) and recovered more quickly (*τ*_r0_= 0.736 s). Table 4 shows full results of the model across these 3 types as well as calculated paired Z-scores (Eq. 6) for P0 and *τ*_r0_. The heterogeneity in layer 2/3 response dynamics made it difficult to constrain the model and thus were not included in this analysis.

## Discussion

We leveraged the sub-millisecond sampling, high gain, and low noise of multipatch recordings to investigate the functional connectivity and short-term dynamics of recurrent synapses in the adult mouse and human cortex. We observed sparse recurrent connections between excitatory neurons in layers 2/3 through 6 in adult mouse visual cortex and layers 2 through 6 of adult human cortex. We supplemented mouse multipatch experiments with high throughput 2P optogenetic stimulation to sample connectivity at greater distances than is generally feasible using the multipatch approach. The large majority of excitatory recurrent connections in mouse cortex exhibited short-term synaptic depression.

Recordings made from *in vitro* slice preparations offer a unique combination of temporal resolution, low resistance intracellular access, and long term stability. Although these features provide an excellent environment for multipatch recordings, there are also associated limitations that must be considered carefully. Estimates of connectivity derived from multipatch experiments in brain slices should be considered as a lower bound on the underlying population connectivity due to sensitivity to false negatives from several sources. These effects may contribute to differences in reported connectivity across studies. A fraction of synaptic connections are expected to be severed during slicing; one estimate of connectivity perturbed by slicing approaches 50% (Levy and Reyes 2012). The effect on measured connection probability depends on the thickness of the slice, the depth of recorded cells from the cut surface, the morphology of recorded cells, and the distance between them. Although we minimize lost connections by patching deep in the slice (>40 µm) and by selecting cells in close proximity, this is still a likely source of false negatives in our data. Another fraction of synapses are expected to be either too weak (Isaac et al 1995) or too distal from the recording pipette to be detected. The magnitude of this effect is difficult to estimate, but our initial analysis hints that our methods are sensitive enough to capture the majority of synapses. To obtain more accurate estimates of connectivity, it will be necessary to combine these results with other methods such as *in vivo* multipatch recordings, transsynaptic tracing, and serial section electron microscopy. These methods are also limited, but in each case the constraints are different and potentially complementary.

There is a wide range of reported rates of recurrent connectivity among excitatory neurons in rodent studies. One suggestion is that differences between the juvenile and adult rodent can explain the variance (Jiang et al. 2016). To avoid changes associated with development, we carried out our experiments in the adult (about two months old) cortex. Nevertheless, our conclusion that recurrent connectivity is sparse but not absent is similar to results from experiments in other adult (Cossell et al. 2015, Lee et al. 2016) and juvenile animals (Mason et al. 1991, Holmgren et al. 2003, Song et al. 2005, Sjostrom et al. 2001, Morishima et al. 2011, Perin et al. 2011, Lefort et al. 2009, Levy and Reyes 2012). A notable difference, however, is that we never observed rates of recurrent connectivity as high or synaptic amplitudes as large as those reported in juvenile rodents, consistent with the observation that the rate of recurrent connectivity and synaptic strength declines with age (Burgeois and Rakic 1993, Reyes and Sackman 1999).

Different rates of recurrent connectivity across neuronal classes may suggest a mechanism for observed differences in their functional influence on downstream targets. In this context, it is interesting that in the mouse, all three classes of deep projection neuron had different rates of recurrent connectivity. This was approximately 10% in Sim1 (layer 5; subcortically projecting) expressing cells, whereas Tlx3 expressing neurons (layer 5; intracortically projecting) interconnect approximately half as frequently (∼4%). Relatively high rates of recurrent connectivity are not a generalizable property of subcortically projecting cells, as Ntsr1 expressing cells (layer 6) had the lowest rate of intralaminar connectivity among all excitatory cell types tested here.

How do the rates of recurrent connectivity in mouse compare to other species? In human cortex, we and others (Molnar et al. 2008) find that the frequency of connectivity among layer 3 excitatory neurons is at least two and half times greater than the highest connectivity rate observed in the adult mouse. It may be that high rates of recurrent connectivity in layer 3 are a circuit feature of higher mammals as this is also reported to be the case in cat and monkey (Kisvarday et al 1986, McGuire et al 1991, Bopp et al 2014). Future work will examine how short-term dynamics and/or inhibitory feedback may counterbalance recurrent excitatory connectivity to prevent runaway excitation.

In our study, the distance dependent connection probability profiles observed in all excitatory types appear to fall off with distance and are consistent with connectivity depending on the extent of overlap between neighboring axons and dendrites (Peters 1979, Binzegger et al. 2004, Braitenberg and Schüz 1998, van Pelt and van Ooyen 2013). It is unlikely that the distance dependent connectivity profiles we observed are an artifact of tissue preparation as it has previously been demonstrated that the truncation of neuronal processes reduces overall connectivity but maintains the spatial pattern of connections (Stepanyants et al. 2009, Levy and Reyes 2012). Additionally, 2P optogenetic experiments allowed for interrogating potential connections at much larger distances. Further effort is needed to determine whether our connectivity profiles can be predicted from neuronal morphology.

Recurrent excitatory synapses most commonly exhibit short-term depression (Markram 1997, Reyes and Sackmann 1999, Lefort and Petersen 2017), which is thought to normalize the synaptic gain across different firing frequencies (Abbott et al. 1997) as well as minimize runaway excitation within recurrent circuit elements. In our sample, the rate of entry into depression varied among Cre types, which suggests differences in initial release probability. In support of this hypothesis, we also observed that Tlx3 expressing neurons exhibited the slowest entry into depression had the smallest initial EPSP amplitudes and a relatively high CV. Nevertheless, the short-term dynamics of the Cre expressing neurons were uniformly depressing, and their maximal depression was also similar. In contrast, layer 2/3 pyramidal cell STP was more evenly distributed between facilitating and depressing. This is in agreement with a previous study by Lefort and Petersen (2017). Although less common, facilitation in recurrent excitatory synapses has also been observed in the medial prefrontal cortex and is hypothesized to play a role in reverberant activity (Wang et al. 2006).

Ultimately, we seek a description of the cortical circuit from which mechanistic computational models can be built and hypotheses about cortical function can be tested. Although many parts of the circuit have been described in the past, incompatibilities between experiments have made it difficult to assemble a complete, coherent picture of the whole. We have taken steps toward ensuring that our results can be interpreted in the context of future experiments, but more work is needed to generate a consistent description of the cortical circuit. To that end, we have begun a large-scale project to replicate these measurements across a wider variety of cell types in the mouse and human cortex; the results of our early-stage data collection presented here suggest that systematic and standardized characterization will provide a detailed, quantitative, and comprehensive description of the circuit wiring diagrams and will facilitate the investigation of circuit computation.

## Methods

### Animals and Tissue Preparation

Adult mice of either sex (mean age P45 ± 4; SD) were housed and sacrificed according to protocols approved by the Institutional Care and Use Committee at the Allen Institute (Seattle, WA), in accordance with the National Institutes of Health guidelines. Transgenic mouse lines were used for experimentation and chosen based on cortical layer specific expression and/or known projection patterns. In the following mouse lines, subpopulations of excitatory neurons are selectively labelled with fluorescent reporters (tdTomato or GFP): Tlx3-Cre_PL56;Ai14 363 (n=57), Sim1-Cre_KJ18;Ai14 (n=20), Rorb-T2A-tTA2;Ai63 (n=28), Ntsr1-Cre_GN220;Ai140 (n=10) (Allen Institute; see also http://connectivity.brain-map.org/transgenic). Two drivers, Sim1-Cre (subcortical projecting; CS or PT type; Allen Brain Atlas, http://connectivity.brain-map.org/) and Tlx3-Cre, (corticocortical projecting; CC or IT type; Kim et al 2015), were used to label layer 5 pyramidal cells, in order to sample projection-specific subpopulations. For optogenetic experiments, Tlx3-Cre driver mice were bred with ROSA26-ZtTA/J mice (Jackson Laboratory) and Ai136 mice (Daigle et. al 2017), in which a fusion of the ReaChR opsin (Lin 2013) with EYFP is expressed from the TIGRE locus (Zeng et al. 2008) in a Cre-and tTA-dependent manner. Previous studies have emphasized the differences in cortical connectivity particularly at older ages. To assess whether age impacted the results reported here, a subset of experiments were repeated for Sim1 connections in older mice (mean age 61 ± 1; SD, n=10). We saw no difference in recurrent connectivity rate (<100 µm; P40-60: 36/423, >P60: 15/269, Fisher’s p = 375 0.23) or response amplitude (P40-60: 0.53±0.12 mV, >P60: 0.59±0.2 mV, p = 0.98 KS test) among across the two time points.

Animals were deeply anesthetized with isoflurane and then transcardially perfused with ice-cold oxygenated artificial cerebrospinal fluid (aCSF) containing (in mM): 98 HCl, 96 N-methyl-d-glucamine (NMDG), 2.5 KCl, 25 D-Glucose, 25 NaHCO3, 17.5 4-(2-hydroxyethyl)-1-piperazineethanesulfonic acid (HEPES), 12 N-acetylcysteine, 10 MgSO4, 5 Na-L-Ascorbate, 3 Myo-inositol, 3 Na Pyruvate, 2 Thiourea, 1.25 NaH2PO4·H2O, 0.5 CaCl2, and 0.01 taurine (aCSF 1). All aCSF solutions were bubbled with carbogen (95% O2; 5% CO2).

Acute parasagittal slices (350 µm) containing primary visual cortex from the right hemisphere were prepared with a Compresstome (Precisionary Instruments) in ice-cold aCSF 1 solution at a slice angle of 17° relative to the sagittal plane in order to preserve pyramidal cell apical dendrites. Slices were then allowed to recover for 10 minutes in a holding chamber (BSK 12, Scientific Systems Design) containing oxygenated aCSF 1 maintained at 34°C (Ting et al. 2014). After recovery, slices were kept in room temperature oxygenated aCSF holding solution (aCSF 2) containing (in mM): 94 NaCl, 25 D-Glucose, 25 NaHCO3, 14 HEPES, 12.3 N-acetylcysteine, 5 Na-L-Ascorbate, 3 Myo-inositol, 3 Na Pyruvate, 2.5 KCl, 2 CaCl2, 2 MgSO4, 2 Thiourea, 1.25 NaH_2_PO_4_ •H_2_O, 0.01 Taurine for a minimum of one hour prior to recording.

Human tissue surgically resected from adult cortex was obtained from patients undergoing neurosurgical procedures for the treatment of symptoms associated with epilepsy or tumor. Data were collected from 67 total slices from 22 surgical cases (17 epilepsy, 5 tumor, mean age 40 ± 17 years; SD). Tissue obtained from surgery was distal to the core pathological tissue and was deemed by the physician not to be of diagnostic value. Specimens were derived from the temporal lobe (13 epilepsy, 4 tumor) and the frontal lobe (4 epilepsy, 1 tumor). Specimens were placed in a sterile container filled with prechilled (2-4°C) aCSF 3 containing decreased sodium replaced with NMDG to reduce oxidative damage (Zhao et al., 2011) composed of (in mM): 92 NMDG, 2.5 KCl, 1.25 NaH_2_PO_4_, 30 N_a_HCO_3_, 20 HEPES, 25 glucose, 2 thiourea, 5 Na-ascorbate, 3 Na-pyruvate, 0.5 CaCl_2_·4H_2_O and 10 MgSO_4_·7H2O. pH was titrated to 7.3–7.4 with HCl and the osmolality was 300-305 mOsmoles/Kg. Surgical specimens were transported (10-40 minutes) from the surgical site to the laboratory while continuously bubbled with carbogen.

Resected human tissue specimens were trimmed to isolate specific regions of interest, and larger specimens were cut into multiple pieces before trimming. Specimens were mounted in order to best preserve intact cortical columns (spanning pial surface to white matter) before being sliced in aCSF 3 using a Compresstome. Slices were then transferred to oxygenated aCSF 3 maintained at 34°C for 10 minutes. Slices were kept in room temperature oxygenated aCSF holding solution (aCSF 4) containing, in mM: 92 NaCl, 30 NaHCO3, 25 D-Glucose, 20 HEPES, 5 Na-L-Ascorbate, 3 Na Pyruvate, 2.5 KCl, 2 CaCl_2_, 2 MgSO_4_, 2 Thiourea, 1.2 NaH_2_PO_4_·H_2_O for a minimum of one hour prior to recording.

### Electrophysiological Recordings

Recording slices from mouse and human tissue were processed in largely the same manner, with a key difference being the external calcium concentration used for recording. Human slices were held in aCSF containing 1.3 mM calcium while mouse slices utilized 2.0 mM calcium. Below we discuss the full preparation for slice processing as well as the rationale for this calcium difference.

Slices were transferred to custom recording chambers perfused (2 mL/min) with aCSF maintained at 31-33°C, pH 7.2-7.3, and 30-50% oxygen saturation (as measured in the recording chamber). aCSF (aCSF 5) contained (in mM), 1.3 or 2 CaCl2 (2.0 in mouse experiments and either 1.3 or 2.0 in human experiments), 12.5 D-Glucose, 1 or 2 MgSO_4_, 1.25 NaH_2_PO_4_·H_2_O, 3 KCl, 18 NaHCO_3_, 126 NaCl, 0.16 Na L-Ascorbate.

The concentration of calcium in the external recording solution affects release probability and other aspects of synaptic dynamics (Borst 2010, Pala & Petersen 2015, Jouhanneau 2015, Urban-Ciecko et al 2015). Although concentrations close to 1 mM are expected to most closely approximate *in vivo*-like synaptic dynamics, most prior multipatch studies used elevated calcium concentrations to increase the strength of synaptic currents and improve throughput. In our mouse recordings, we used 2.0 mM CaCl2 to be consistent with previous connectivity studies (Markram et. al 1997, Reyes and Sakmann 1999, Perin et. al 2011, Jiang et. al 2015) and to help ensure the success of our system integration test. In a limited test among recurrent Sim1 connections, the difference in synapse strength was statistically insignificant between 2.0 mM and 1.3 mM CaCl_2_ (2.0mM CaCl_2_ single-pulse amplitude = 501 ± 108 µV, n=9; 1.3mM CaCl_2_ = 383 ± 90 µV, n=15; KS test p = 0.47). We selected 1.3 mM [Ca++]_e_ for our human experiments because of reports that the synaptic strength is higher than in mouse and to minimize the complex events that can be initiated by individual spikes in human tissue (Molnar et al. 2008) that make identifying monosynaptic connectivity challenging. Future mouse recordings will also be carried out at 1.3 mM [Ca++]_e_ in order to capture the physiological synaptic dynamics.

Slices were visualized using oblique infrared illumination using 40x or 4x objectives (Olympus) on a custom motorized stage (Scientifica), and images were captured using a digital sCMOS camera (Hamamatsu). Pipette positioning, imaging, and subsequent image analysis were performed using the python platform acq4 (acq4.org, Campagnola et al. 2014). Eight electrode headstages (Axon Instruments) were arranged around the recording chamber, fitted with custom headstage shields to reduce crosstalk artifacts, and independently controlled using triple axis motors (Scientifica). Signals were amplified using Multiclamp 700B amplifiers (Molecular Devices) and digitized at 50-200kHz using ITC-1600 digitizers (Heka). Pipette pressure was controlled using electro pneumatic pressure control valves (Proportion-Air) and manually applied mouth pressure.

Recording pipettes were pulled from thick-walled filamented borosilicate glass (Sutter Instruments) using a DMZ Zeitz-Puller (Zeitz) to a tip resistance of 3-8 MΩ (diameter ∼1.25 µm), and filled with internal solution containining (in mM): 130 K-gluconate, 10 HEPES, 0.3 ethylene glycol-bis(β-aminoethyl ether)-N,N,N’,N’-tetraacetic acid (EGTA), 3 KCl, 0.23 Na2GTP, 6.35 Na2Phosphocreatine, 3.4 Mg-ATP, 13.4 Biocytin, and either 25 µM Alexa-594 (for optogenetic experiments), 50 μM Cascade Blue dye, or 50 µM Alexa-488 (osmolarity between 280 and 295 mOsm titrated with ∼4 mM sucrose, pH between 7.2 and 7.3 titrated with ∼10 mM KOH). The liquid junction potential between our internal solution and aCSF 5 was measured to be 9.40 mV ± 0.59; SD. All electrophysiological values are reported without junction potential correction.

In experiments on human tissue and wild-type mice, clusters of up to 8 excitatory neurons (depth from slice surface ≥ 40 µm) were selected based on cortical layer and somatic appearance. In transgenic mice, cells were also targeted based on fluorescent reporter expression. All cells were confirmed as excitatory post-experiment either by their synaptic currents onto other recorded neurons (Fig S1A) or by their pyramidal morphology, visualized using either biocytin (Fig 1A) or fluorescent dye from the pipette (Fig 1B). Cell intrinsic fluorescence was confirmed post-hoc via manual inspection of image stacks to evaluate signal overlap of the transgenic fluorescent reporter and the fluorescent dye introduced via pipettes (Fig 1B). Whole-cell patch clamp electrophysiological recordings were performed at −70 mV to preferentially measure excitatory inputs. Custom software, Multi-channel Igor Electrophysiology Suite, written in Igor Pro (WaveMetrics), was used for data acquisition and pipette pressure regulation. A brief, 10 ms test pulse was used to monitor access and input resistance over the duration of the recording. Resting membrane potential was maintained within 2 mV using automated bias current injection during the inter-trial interval. During recordings, cells were stimulated using brief current injections (1.5 or 3 ms) to drive trains of 12 action potentials (Figure S1A) at frequencies of 10, 20, 50, or 100 Hz to induce short-term plasticity (STP). A delay period inserted between the 8th and 9th pulses allowed testing of recovery from STP. In most recordings this delay period was 250 ms; for 50 Hz stimulation, longer delay periods (500, 1000, 2000, and 4000 ms) were used as well (see Fig 5A). Stimuli were interleaved between cells such that only one cell was spiking at a time, and no two cells were ever evoked to spike within 150 ms of each other.

### Data Analysis

Postsynaptic recording traces were aligned to the time of the presynaptic spike evoked from the stimuli described above (Figure S1B). Postsynaptic potentials (PSPs) were identified by manual inspection of individual and averaged pulse-response trials. A classifier (described below) was later used to highlight possible identification errors, which were then manually corrected. Connection probabilities within 100 µm intersomatic distance were compared between cell types using Fisher’s exact test of 2×2 contingency tables (connected, unconnected). Bonferroni method was used to report corrected p values for multiple comparisons among cell types. The relationship between connectivity and intersomatic distance (measured from 3D cell positions) was analyzed by binning connections in 40 µm windows and calculating the 95% Jeffreys Bayesian confidence interval for each bin.

Subsets of the 97 mouse and 57 human connections found in this study were analyzed for strength, kinetics, and STP based on specific quality control criteria (Fig S1C-G, Table 1). EPSP strength, kinetics, and coefficient of variance (CV) measurements (Fig 1 and 3) were conducted on the first pulse response of 10, 20, and 50 Hz stimulation trains which were time-aligned to the presynaptic spike and averaged for each connection. Connections were included for strength and kinetics analysis according to the analysis flowchart in Fig S1E and F. Briefly, the postsynaptic cell had an auto bias current less than 800 pA (mean bias current −176 ± 213 pA), there was no spontaneous spiking, the stimulus artifact was minimal (<30 µV), and the PSP was positive. Individual recording sweeps were included if the baseline potential drift was smaller than ± 5 mV from holding (−70 mV) and the mean baseline 10 ms preceding stimulation was less than 3 standard deviations of the mean baseline across sweeps. In the QC passed data, strength and kinetics were measured from a double exponential fit that approximates the shape of the PSP:

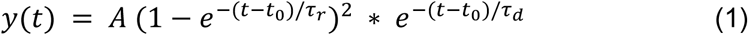

Best fit parameters were obtained using the Non-Linear Least-Squares Minimization and Curve-Fitting package for Python (LMFIT; Newville et al., 2014). To improve the quality of fitting, the root mean square error was weighted (WRMSE) differently throughout the trace. The rising phase of the PSP was most heavily weighted, the baseline and decay regions were intermediately weighted, and the region of the presynaptic stimulus, which often contained crosstalk artifacts was masked. Amplitude was measured as the peak of the PSP fit (Fig S1B). Kinetics were measured from connections in which the WRMSE of the fit was less than 8. Latency is reported as the duration from the point of maximum *dV/dt* in the presynaptic spike until the foot of the PSP (Fig S1B), taken from the x-offset in the double exponential fit. Rise time is reported as the duration from 20% of the peak until 80% of the peak of the PSP (Fig S1B). Significance of differences in PSP amplitude, latency, and rise time across layers or Cre-types were assessed with a Kruskal-Wallis test.

STP (Fig 5 and 7) was measured from a similar subset of connections that included the quality control criteria above and also excluded responses smaller than 0.5 mV in amplitude to minimize the effect of noise on mean response which might impact the model (Fig S1G, Eq. 4 and 5). Connections or individual sweeps that had a baseline holding potential of −55 mV (± 5 mV) were reintroduced for this analysis if they met the QC criteria. Normalized PSP amplitudes (relative to the first pulse) were estimated using an exponential deconvolution (□=15 s; Richardson et al., 2008) to compensate for summation from prior PSPs and to increase signal-to-noise in measuring PSP amplitudes:

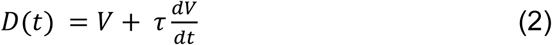

Although the fixed deconvolution time constant of 15 ms may differ significantly from the actual time constant of each cell, in practice this has little effect on the normalized amplitudes used in STP measurements (for example, using this time constant to measure amplitudes from a simulated 100Hz train with a cell time constant of 30 ms only resulted in 3% error in the measurement of PSP amplitudes relative to the first pulse; data not shown). The peak amplitudes from the deconvolved traces were used to measure the change in response magnitude over the course of stimulus trains. We measured the magnitude of short-term depression or facilitation using the ratio between the first and last (eighth) pulses in an induction pulse train, whereas recovery from depression or facilitation was measured by the ratio between the first pulse and the ninth pulse, which followed a recovery delay. Kruskal-Wallis tests were used to assess significance of STP between multiple layers. A descriptive model was used to capture features of short-term depression in Rorb, Sim1, and Tlx3 connections (Eq. 4 and 5).

### Automatic synapse detection

To aid in the detection of synaptic connections, a support vector machine classifier (implemented with the “sklearn” python package, Pedregosa et al. 2011) was trained to discriminate between experiments in which synaptic currents were either visible or not visible to a human annotator. The classifier required a diverse set of features that were pre-processed from the raw pulse response recordings. Averaged pulse responses were characterized by curve fitting (Eq. 1; Fig. 1B) and the fit parameters as well as the normalized RMS error were provided as features to the classifier. Additionally, individual pulse response recordings were analyzed by measuring the amplitude and time of the peak of each exponentially deconvolved response over a 3 ms window, compared to a 10 ms window preceding the stimulus pulse (Fig. 1B, bottom). Although these individual measurements were often noisy (average background RMS noise 607±419 µV), their distribution over hundreds of trials could be compared to similar distributions measured from background noise (e.g. Fig. 4C). Distributions were compared using a Kolmogorov-Smirnov test (from the “scipy.stats” Python package) and the p values were used as input features for the classifier.

After training on 1854 manually labeled examples, the classifier was tested against a withheld set of 2642 examples and achieved an overall accuracy of 95% (56/61 true positive, connected; 2457/2581 true negative, not connected). False positives and negatives were manually reassessed and frequently found to have been misclassified during the initial manual annotation; such instances were corrected for in measurements of connectivity.

### Analysis of synapse detection sensitivity

To measure the minimum detectable PSP size for each connection probed, artificial PSPs were added to recordings of background noise taken from the postsynaptic cell. PSPs were generated using Equation 1 with a foot-to-peak rise time of 2 ms (except where specified in Fig 3D). PSP latencies were selected from a gaussian distribution centered at 2 ms with a 200 µs standard deviation. PSP amplitudes were generated by the product of two random variables: one binomially distributed (p=0.2, n=24) to mimic stochastic vesicle release, and the other normally distributed (mean=1, SD=0.3) to account for differences in vesicle size and receptor efficacy. PSPs were then scaled uniformly to achieve a specific mean amplitude. The resulting simulated responses were qualitatively similar to typical synaptic responses encountered in our dataset, although they lacked the synapse-to-synapse variability in CV, due to the selection of fixed distribution parameters listed above.

For each connection probed, the number of simulated PSPs generated was the same as the number of presynaptic spikes elicited during the experiment. These PSPs were then fed through the same preprocessing and classification system that was used to detect synaptic connections in real data, and the classification probability was calculated from the classifier (using sklearn.svm.SVC.predict_proba). This process was repeated 8 times (with PSPs generated randomly each time) and the average classification probability was recorded. This yields an estimate of whether a synaptic connection would be detected or overlooked, given the combination of sample count, background noise characteristics, and PSP strength and kinetics.

By repeating this process for several different values of mean PSP amplitude, we could identify, for each putative connection probed, a plausible minimum detectable PSP amplitude. This minimum detectable amplitude was defined as the PSP amplitude at which the classifier would detect the synapse in 50% of trials (interpolated from adjacent amplitudes).

### PSP amplitude run-down over duration of experiment

The amplitude of the PSPs initiated by the first pulse of the stimulus trains was characterized over time for the same subset of connections described for the strength and kinetics analysis. In order to discount variations in the measurements of individual first pulses, the run-down was characterized via a linear regression of PSP amplitude versus time. Here, the total run-down is reported as the percentage decrease in amplitude of a PSP at the beginning and end of an experiment (average duration was 24.4 ± 10.8 minutes) specified by the linear regression, i.e. percent run-down,

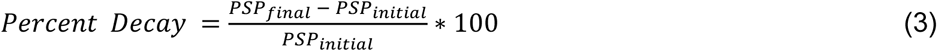

All synapses where the beginning and ending PSP amplitude straddled zero, which can happen for the weaker synapses, were excluded from analysis as they produce aberrant percentage values. On median, the following overall run-downs were observed: layer 2/3 to layer 2/3 connections 11%, Rorb to Rorb 31%, Sim1 to Sim1 22%, Tlx3 to Tlx3 6%.

### Theoretical Synaptic Modelling

Synaptic depression was modelled via depletion of vesicles (Hennig, 2013; Mongillo et al., 2008; Richardson et al., 2005),

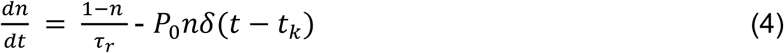

where *n* is the fraction of available vesicles, *P*_0_ is the release probability, *t*_k_ is the time of presynaptic spike and *τ*_r_ is the time constant for vesicle replenishment. The speed of replenishment can vary over time depending on the history of presynaptic spikes, which can be captured by time constant *τ*r evolving according to Eq. 5 (Fuhrmann et al., 2002; Hennig, 2013),

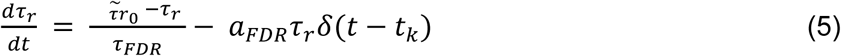

where *τ*_*FDR*_ is the time constant of use-dependent replenishment, *a*_*FDR*_ represents the amount of updates elicited by a presynaptic spike and 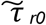 is the baseline time constant. For Rorb, Sim1, and Tlx3 synapses, we optimized the parameters (P0, 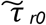, *τ*_*FDR*_ and *a*_*FDR*_) to account for time courses of PSPs. Specifically, PSPs were averaged over all available synapses depending on stimulation frequencies and delays between 8^th^ and 9^th^ presynaptic pulses and fitted to the model. We used LMFIT (Newville et al., 2014) to perform non-linear least-square minimization.

Paired Z-scores for P_0_ and *τ*_*r0*_ were calculated from the standard error returned during parameter optimization according to Eq 6.,

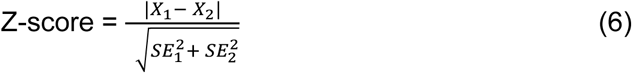

Where X_1_ and X_2_ are P_0_ or *τ* _*r0*_ for 2 groups and their associated standard error

### Histology and Morphology

After completing electrophysiological recordings, slices were transferred from the recording chamber and fixed in solution containing 4% PFA and 2.5% glutaraldehyde for 2 days (>40 hours) at 4°C. After fixation, slices were transferred and washed in phosphate buffer saline (PBS) solution for 1-7 days.

Sections were processed using 3,3’-diaminobenzidine (DAB) peroxidase substrate kit to identify recorded neurons filled with biocytin. Free floating sections were first incubated with 5 µM 4’,6-diamidino-2-phenylindole (DAPI) in PBS for 15 minutes at room temperature and then triple washed in PBS (3 ×; 10 minutes). Sections were transferred to a 1% H2O2 (in PBS) for 30 minutes and then triple washed in PBS. A DAB substrate kit (VectorLabs) was used to stain for neurons filled with biocytin. Sections were mounted on gelatin-coated slides and coverslipped with Aqua-Poly/Mount (Polysciences).

Slides were imaged on an AxioImager Z2 microscope (Zeiss) equipped with an Axiocam 506 camera (Zeiss) and acquired via the Zeiss Efficient Navigation software. Tiled mosaic images of whole slices were acquired via automated scanning and stitching of several 20X images to generate both brightfield and DAPI images of the entire slice.

### Two photon optogenetic experiments

Connectivity mapping experiments were performed on a two-photon laser scanning microscope (Bruker Corp) with a tunable pulsed Ti:Sapphire laser (Chameleon Ultra, Coherent) for imaging, and a fixed wavelength (1060 nm) pulsed laser (Fidelity Femtosecond, Coherent) for stimulation. A 63x, 1.0 NA water immersion objective (Zeiss) was used for all experiments. Two-photon images were acquired with PrairieView software (Bruker Corp), and stimulation targets were manually placed on these reference images to target ReaChR-positive cells. Photoactivation stimuli were triggered by a TTL pulse generated within MIES acquisition software. The voltage output controlling the photoactivation Pockels cell was recorded within MIES for post-hoc alignment of physiological recordings with the timing of photoactivation. To characterize the effectiveness and specificity of stimulation parameters, we made loose seal recordings on to EYFP/ReaChR-labelled neurons (Figure S2A). For all data presented here, the photostimulation pattern consisted of a spiral 5 μm in diameter with 5 revolutions traced over a 25 ms duration. We first determined the minimum light power necessary to evoke reliable firing of action potentials. This minimum power varied across cells from 2.6 −20.3 mW (Figure S2B). A photo-stimulus of 18 mW intensity was sufficient to evoke spiking in 92% of cells tested (12/13 cells). The average latency of firing at this power was 12.9 ± 6.1 ms and the associated jitter was 0.98 ± 0.58 ms (Figure S2C,D).

Within the same experiments, we characterized the spatial specificity of these stimulation parameters. First, to determine the probability of off-target photoactivation of cells within the same focal plane, we delivered stimuli in a radial grid pattern containing 7 spokes with stimuli spaced 10, 20 and 30 µm away from the center of the recorded cell (Figure S2E). Spike probability fell to 0.5 at a lateral distance of 12.0 µm. Finally, we determined the axial resolution of our photoactivation paradigm by offsetting the focus of the objective relative to the recorded cell. Consistent with previous studies, (Packer et al., 2012; Prakash et al., 2012) axial resolution was inferior compared to lateral resolution (spike probability = 0.5 at 26.7 µm) but was still near 656 cellular resolution (Figure S2F).

For two photon mapping experiments, 1-2 neurons were patched and membrane potential was maintained near −70 mV with auto bias current injection. Neurons were filled with 50 µM Alexa-594 to visualize cell morphology (Figure S3A). The orientation of the apical dendrite was utilized to align photostimulation sites across experiments in downstream analyses. Each putative presynaptic neuron was stimulated 10-20 times, with the parameters described above. Photo stimulation was performed in ‘rounds’ during which EYFP-labelled neurons within a single field of view were sequentially targeted (3-12 neurons/round). Stimulation protocols were constrained such that the inter-stimulus interval between neurons was ≥ 2 s and the inter-stimulus interval for a given neuron was ≥ 10 s.

Photostimulus responses were scored as connection, no connection or as containing a direct stimulation artifact by manual annotation. To assist in these user-generated calls, we incorporated a signal-to-noise measure for our optogenetic mapping data. Current clamp traces were low pass filtered at 1 kHz and baseline subtracted. The voltage-deconvolution technique (Eq. 2) was then applied. The value of t was set between 10-40 ms. Deconvolved traces were 671 high pass filtered at 30 Hz, and peaks larger than 3 standard deviations above pre-stimulus baseline were used for further analysis (Fig 4 S3B,C). We then measured the number of peaks in both ‘signal’ and ‘noise’ regions. The ‘signal’ region was a 100 ms window 5-105 ms after the 674 onset of the photostimulus, and the ‘noise’ region was a 100 ms window 145-45 ms before the stimulus onset. To compensate for jitter known to be present in two-photon mediated stimulation, we determined a 10 ms subset within each 100 ms window that gave the maximum number of unique trials containing threshold-crossing events. The median of the peak within this 10 ms window was found across all trials in both ‘signal’ and ‘noise’ regions, and the mean of a 25 ms window preceding both regions was subtracted to produce our final signal and noise values.

We plotted the signal against the noise for all stimulus locations (Fig 4 S3D), and found that most points with a high signal-to-noise ratio contained either a synaptic response or an artifact produced by direct stimulation of the recorded (opsin-expressing) cell. 93% (13/14) manually identified connections had a signal to noise ratio > 1.5 (Fig 4 S3E). By contrast, the same was true of only 1.7% (24/1416) of cells scored as ‘no connection’. Therefore, our signal to noise analyses highlight quantitatively distinct features of our connection calls. The presence of direct stimulation artifacts prevents us from unambiguously identifying synaptic connections between nearby neurons. Therefore, when estimating connection probability by two-photon optogenetics, we did not include putative presynaptic cells within 50 µm of the recorded neuron where the direct stimulation artifact was largest (Fig 4F).

## Acknowledgments

The authors thank the Allen Institute founder, Paul G. Allen, for his vision, encouragement and support. Thank you to the Allen Institute for Brain Science Tissue Processing, Histology, Imaging, and Morphology and 3D reconstruction teams for preparing and processing mouse tissue. We further thank the Tissue Procurement, Tissue Processing, and Facilities teams for help in coordinating the logistics of human surgical tissue collection, transport, and processing. We are also grateful to our collaborators at the local hospital sites, including Tracie Granger, Caryl Tongco, Matt Ormond, Jae-Guen Yoon, Nathan Hansen, Niki Ellington, Rachel Iverson (Swedish Medical Center), Carolyn Bea, Gina DeNoble and Allison Beller (Harborview Medical Center). We thank Dirk C. Keene at Harborview/UW for consultation and support. This work was supported by the Allen Institute for Brain Science, National Institutes of Health grant U01MH105982 to H.Z., and the Howard Hughes Medical Institute grant to GJM.

## Supplement

**Figure S1.**
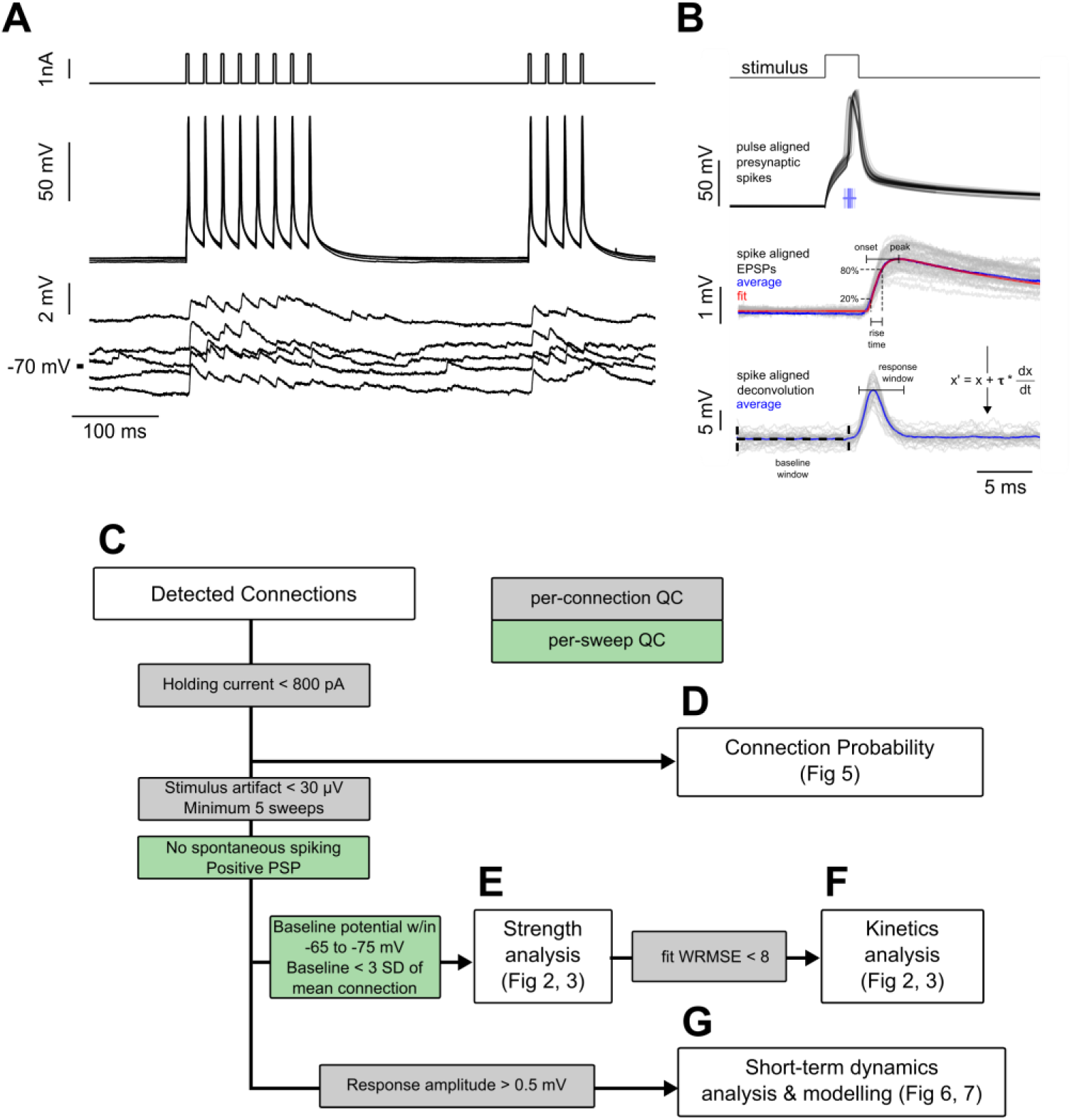
Experiment methodology and analysis workflow. **A.** Example connected pair showing the stimulation pulses (top) and action potentials (middle) in the presynaptic cell; monosynaptically evoked EPSPs (bottom) in the postsynaptic cell. Traces represent every fifth sweep from the 50 Hz protocol used to measure recovery from STP at a delay of 250 ms. **B.** Following repeated stimulation, the response to the first spikei n each train of current pulsess was used for EPSP feature analysis. Spikes are shown aligned to the pulse time to illustrate jitter in spike timing. Spike time was defined as the region of maximum dV/dt in the spike trace, as shown in the raster plot corresponding to spike timing of individual spikes. Below, EPSPs are aligned to the spike time prior to fitting the average EPSP (see Equation 1; individual sweeps in grey, average response in blue, fit shown in red). The rise time was calculated as the interval between 20% and 80% of the peak amplitude of the fit. Spike-aligned EPSPs were deconvolved (see Equation 2, shown in figure), and the peak amplitudes of the deconvolved traces were used to measure changes in response amplitude of the course of a spike train. Responses were corrected to the baseline by subtracting the mode of the region between 10 ms and 50 µs prior to stimulus onset (baseline window). Responses were measured as the peak response during a 4 ms window beginning 1 ms after the spike time (response window shown is aligned to mean spike time). **C-G.** Subsets of total connectivity data were used in subsequent analysis. Flowchart shows sweep (green) and connection (grey) level inclusion criteria for data included in each figure. See Table 1 for total number of cells in each criterion.

**Figure S2.**
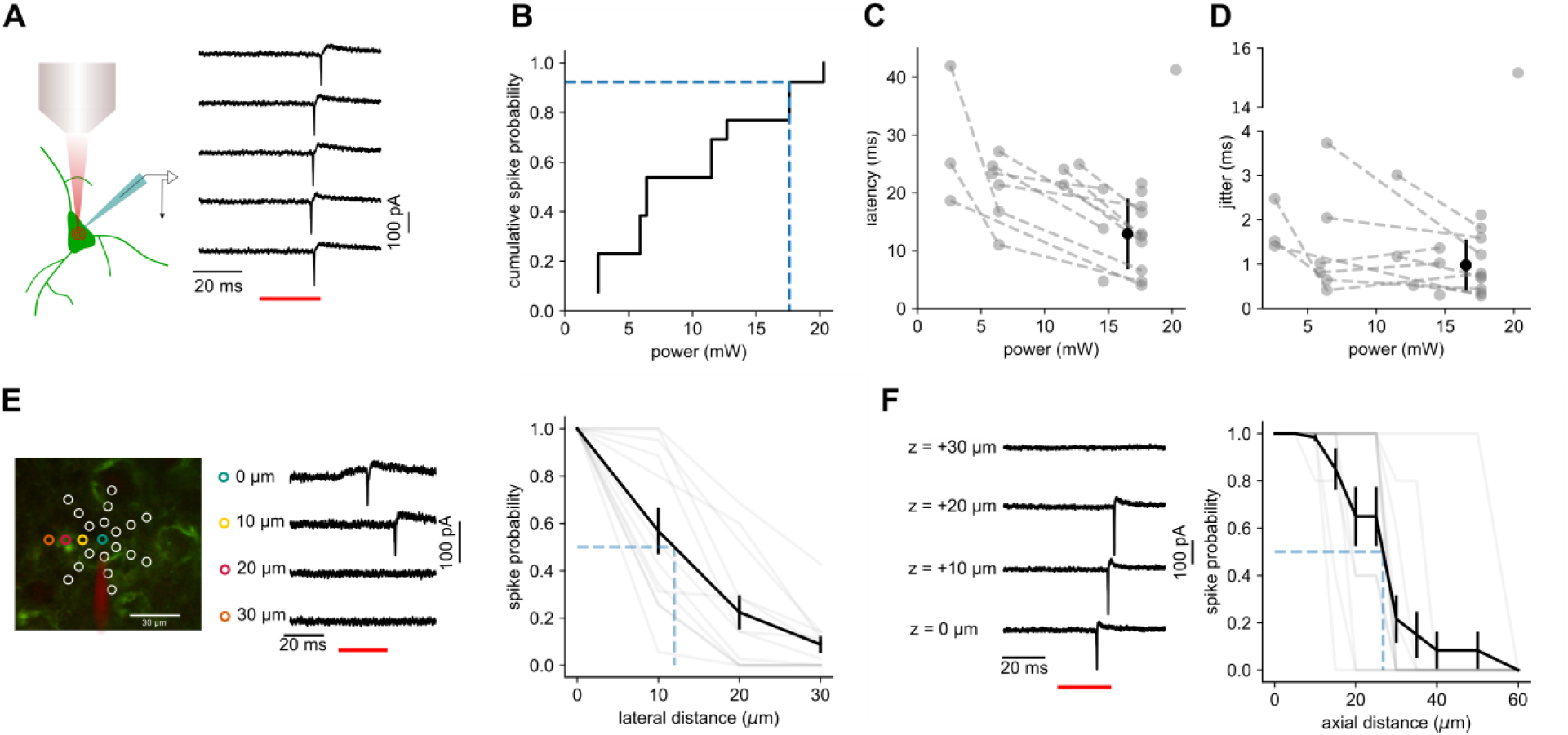
Characterization of two-photon photostimulation. **A.** Cartoon illustrating loose-seal recording configuration utilized to test photostimulation parameters. Example recording of repeated photostimulation or a ReaChR-positive cell. **B.** Cumulative probability plot of minimum power necessary to reliably evoke action potentials for 13 cells. Blue dashed lines indicate light power utilized in mapping experiments and the fraction of cells reliably activated. **C.** Average latency of light-evoked action potentials plotted against photostimulation intensity for individual neurons (grey dashed lines). Filled black circle and error bars represent the mean and standard deviation of latency measured across all cells at a power used for mapping. **D.** Average jitter of light-evoked action potentials plotted against photostimulation intensity. Data from individual cells and population average plotted as in panel C. **E.** *Left:* Example experiment illustrating the radial grid pattern used to measure the lateral resolution of photostimulation and example traces recorded during the photostimulation at indicated locations. *Right:* Probability of generating light-evoked action potentials plotted against lateral distance from the center of the cell. **F.** *Left:* example of responses resulting from photostimulation at indicated axial offsets. *Right:* Probability of generating light-evoked action potentials plotted against axial distance from the center of the cell.

**Figure S3.**
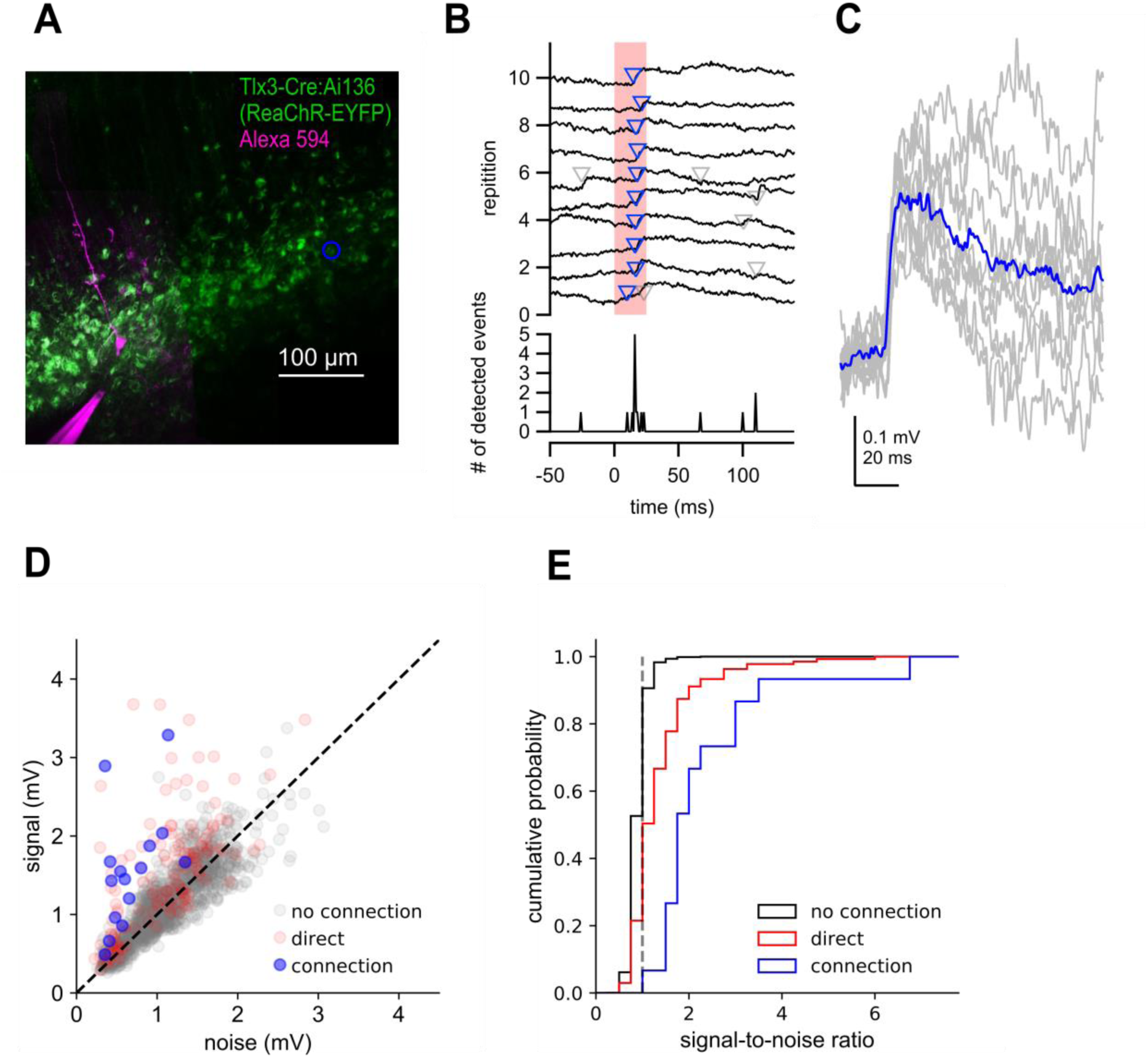
Two-photon optogenetic mapping details. **A.** Maximum intensity projection of Tlx3-Cre:Ai136 slice and a recorded neuron. Blue circle denotes location of stimulated presynaptic neuron. **B.** *Top:* Electrophysiological recording of postsynaptic response to 10 photostimulations of the presynaptic neuron in panel A. Timing of photostimulation indicated by pink shading. Synaptic events detected by exponential deconvolution are indicated by inverted triangles. Events used to produce average synaptic response are show in blue. *Bottom:* Peri-stimulus event histogram. **C.** Individual events aligned by the timing of event detection (grey) and average EPSP (blue). **D.** Signal versus noise plot for all optogenetically-probed presynaptic neurons. **E.** Cumulative probability plot of signal-to-noise ratios for stimulus trials scored as no connection, connection, or containing a direct stimulation artifact. Dashed grey line indicates signal-to-noise ratio = 1.

